# Beyond Brownian motion and the Ornstein-Uhlenbeck process: Stochastic diffusion models for the evolution of quantitative characters

**DOI:** 10.1101/067363

**Authors:** Simon Phillip Blomberg

## Abstract

Gaussian processes such as Brownian motion and the Ornstein-Uhlenbeck process have been popular models for the evolution of quantitative traits and are widely used in phylogenetic comparative methods. However, they have drawbacks which limit their utility. Here I describe new, non-Gaussian stochastic differential equation (diffusion) models of quantitative trait evolution. I present general methods for deriving new diffusion models, and discuss possible schemes for fitting non-Gaussian evolutionary models to trait data. The theory of stochastic processes provides a mathematical framework for understanding the properties of current, new and future phylogenetic comparative methods. Attention to the mathematical details of models of trait evolution and diversification may help avoid some pitfalls when using stochastic processes to model macroevolution.

## Introduction

> “Brownian motion is a poor model, and so is Ornstein-Uhlenbeck, but just as democracy is the worst method of organizing a society “except for all the others”, so these two models are all we’ve really got that is tractable. Critics will be admitted to the event, but only if they carry with them another tractable model.” - J. Felsenstein, r-sig-phylo email list, 8th April 2008.

The parametric estimation of phylogenies depends on having an appropriate model of character evolution (Posada and Crandal, 2001). Molecular systematists are spoiled for choice in this regard. For example, the program jModelTest2 can fit 1624 models of DNA sequence evolution (Darriba et al., 2012). The situation for the comparative analysis of continuous traits is quite different. Here, we have mainly two analytical models in popular use: Brownian motion (BM) and the Ornstein-Uhlenbeck (OU) process. Other models such as “early burst” are also sometimes used (e.g. Blomberg et al., 2003; Ingram et al., 2012). Boucher and Demery (2016) develop a BM model with hard bounds. There have been several extensions to the OU model (see below). There are other approaches to phylogenetic comparative analyses that do not use explicit models of evolution (in terms of being able to write down the appropriate equations). Some nonanalytical models can be used to estimate sampling distributions for regression parameters using computer simulation (Garland et al., 1993), and the evolutionary model for continuous traits can also be altered by applying branch-length transformations (e.g. Blomberg et al., 2003; Freckleton et al., 2002; Grafen, 1989; Pagel, 1999). Lynch (1991) introduced an approach to phylogenetic comparative analyses based on quantitative genetics (see also Hadfield, 2010). I do not consider these approaches to phylogenetic comparative analyses here. Instead, I focus on providing an approach to comparative analyses based on the theory of stochastic processes, which unites BM, OU and other processes in a common statistical and probabilistic framework.

Beginning with the work of Bachelier (1900), a prominent application of stochastic processes has been in finance where models have been developed for stock prices, derivatives, options and other financial products. In that domain, the model of Black and Scholes (1973) has been particularly successful (in terms of citations, if not profits), but research into the theory of stochastic processes is still thriving across a wide range of disciplines, especially the physical sciences (e.g. Einstein, 1956; Freund and Poschel, 2000; Gardiner, 2009; Uhlenbeck and Ornstein, 1930). Although diffusion models are common in epidemiology and other life sciences (Fuchs, 2013), applications in evolutionary biology are rare. The Wright-Fisher model and the Moran model (with various extensions) in population genetics are well-known exceptions (Ewens, 2004; Feller, 1951; Fisher, 1922; Moran, 1958; Wright, 1931). Population geneticists have used these stochastic processes to model microevolution. Here we examine the possible uses of stochastic processes in studies of macroevolution, i.e. evolution above the species level (Benton, 2015; Rensch, 1959; Serrelli and Gontier, 2015; Simpson, 1953; Stanley, 1975), with the aim to provide new models and methods for the phylogenetic comparative analysis of non-Gaussian traits. Such models are necessary because current evolutionary models for quantitative traits can have poor performance (Pennell et al., 2015).

As a way forward, I do not incorporate genetics into any of the following macroevolutionary models: I assume the “phenotypic gambit” (Grafen, 1984). This is a necessary assumption as we almost never have information on the genetic architecture from the fossil record (but see Schraiber et al., 2016). Additionally, the diversity of life suggests that evolutionary constraints on evolution are relatively weak on the deep timescale of tens of millions of years and across many speciation events. Nevertheless, if modern macroevolutionary models are at odds with the established facts of genetics and microevolution, the macroevolutionary models should be discarded. There is some evidence that genetic constraints do not play a strong role in slowing the rate of adaptation (Agrawal and Stinchcombe, 2009). However, Schluter (1996) found evidence for the role of genetic constraints over timespans of the order of ∼ 4 Myr. Analyses of a large dataset of bodysizes, combining paleontological, phylogenetic comparative datasets of extant taxa, and historical studies across many taxa have concluded that on relatively short timescales (< 1 Myr), evolution appears to be bounded; over longer timespans evolution is strongly divergent (Estes and Arnold, 2007; Uyeda et al., 2011). Arnold (1992) provided a synthesis of our knowledge of evolutionary constraints. At that time, there was little evidence for the persistance of constraints over deep time and subsequent reviews, concentrating on the quantitative genetic approach to constraints *via* the G-matrix, have not shed much light on this issue (Blows and Hoffmann, 2005; Futuyma, 2010; Pigliucci, 2007). It seems that while a role for genetic constraints on macroevolutionary timescales cannot be ruled out, there is little evidence and what evidence there is is contradictory.

### Mathematical Background

In order to fully understand the mathematics of stochastic processes, some background is required. At least, some knowledge of Riemann-Stieltjes integrals, as well as some understanding of measure-theoretic probability theory is necessary. Introductory books such as Øksendal (2007) or Klebaner (2012) can be helpful. Gardiner (2009) provides an excellent practical approach which largely ignores the measure-theoretic foundations, but concentrates mainly on applications in the physical sciences. For many purposes, one can ignore the measure-theoretic probability foundations of stochastic processes. However, I urge that some basic grasp of the concepts of probability is desirable in order to understand where the correspondences lie between probability theory and evolutionary theory, where evolutionary interpretations of concepts in probability theory are justifiable, and importantly, where the structure of probability theory as a basis for macroevolutionary theory may break down.

### Brownian Motion

Brownian motion (BM) is named for the movement of pollen grains suspended in water, as first observed by the botanist Robert Brown in 1837, but it is observed in many other multi-particle settings. The mathematics of BM were first analysed by Bachelier (1900), who anticipated almost all the mathematical results of Einstein’s work in 1905 in the context of molecular movement (see Einstein, 1956). Wiener (1923) was the first to rigorously characterise BM as a stochastic process, and hence BM is sometimes also known as the Wiener process. BM was introduced as a model of gene frequency evolution for phylogeny estimation by Edwards and Cavalli-Sforza (1964), and as a model of quantitative character evolution for phylogeny estimation by Felsenstein (1973), who also introduced this model into phylogenetic comparative regression analyses (Felsenstein, 1985).

Let *B*(*t*) be the trait value of a BM process at time *t*. BM has the following defining properties (e.g. Klebaner, 2012). BM has independent increments. *B*(*t*) − *B*(*s*) for *t* > *s* is independent of the past *B*(*u*) where 0 ≤ *u ≤ s*. The increments are also Gaussian. *B*(*t*) − *B*(*s*) has a standard Normal (Gaussian) distribution with a mean *μ* = 0 and variance equal to *t* − *s*. This means we can use all the powerful mathematical machinery appropriate to Gaussian distributions.

Further, the sample paths of a BM process *B*(*t*) have the following properties, for almost every sample path (i.e. other than those of Lebesgue measure zero): *B*(*t*) is a continuous function of *t*. Hence, BM can be used to model continuous traits in continuous time. *B*(*t*) is not monotone in any time interval, no matter how small the interval. BM paths are jagged at all time scales. Despite being continuous, *B*(*t*) is nowhere differentiable. This property makes it difficult to estimate rates of evolution from sample paths, although *σ*^2^ is usually associated with the rate of evolution. The quadratic variation of *B*(*t*) = *t*. That is, the variance of *B*(*t*) increases linearly with *t*. There doesn’t seem to be any biological reason why the variance of a trait should increase linearly with time. Further, this property implies that there are no bounds to evolution and that traits have no physical limits. This is unlikely to be true for any trait (e.g. McGhee, 2015) but see Conway Morris et al. (2015); Vermeij(2015, *ibid.*). More complicated models such as that of Boucher and Demery (1016) impose limits on the variance of sample paths.

BM is useful as a simple model of trait evolution. Its simplicity is due to the above properties, as well as to the fact that it has the Markov property. Further, BM is a martingale, which means that the expectation of the process at time *t* + s is the value of the process at time *t*. That is, *E*(*B*(*t* + *s*)|*ℱ*_*t*_) = *B*(*t*). The Markov and martingale properties simplify the mathematics of working with BM processes. BM lends itself to two evolutionary interpretations. Either it is a model implying no selection and evolution occurs just by random drift, or it can be viewed as a model of very strong selection in a randomly varying environment (see Hansen and Martins, 1996). These interpretations cannot be simultaneously correct, and both are likely to be wrong for any real quantitative trait.

The use of simple BM models for the analysis of comparative data from extant species begins with Felsenstein’s 1985 Phylogenetically Independent Contrasts method. All Ito diffusions use BM as a building block, so BM is included in most quantitative evolutionary models. Felsenstein (2012) has developed a comparative method that allows for the BM evolution of latent traits.

### The Ornstein-Uhlenbeck Process

The OU process was introduced as an improved model for physical Brownian motion, which incorporates the effect of friction (Uhlenbeck and Ornstein, 1930). It also has a long history in evolutionary biology. It can be derived from the consideration of stabilising selection and genetic drift (Lande, 1976). Its use in phylogenetic comparative methods has been promoted by many authors (Beaulieu et al., 2012; Butler and King, 2004; Felsenstein, 1988; Hansen, 1997; Hansen and Martins, 1996; Hansen et al., 2008; Martins and Hansen, 1997). It has the following form:

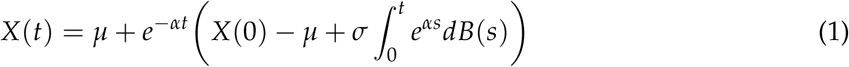

where *μ* is the attracting constant, and the mean of the process at stationarity. Note that *X*(*t*) in (1) depends on *B*(*s*). That is, BM is one building block of the OU process. The biological interpretation of the OU process is controversial. Most authors have interpreted *α* as the strength of a restraining force corresponding to stabilizing selection, and the sample paths as trajectories of evolution of organisms’ traits (e.g. Beaulieu et al., 2012; Butler and King, 2004). However, Hansen (1997); Hansen et al. (2008) interpret the sample paths as paths of an evolutionary optimum itself, subject to an overall central tendency with strength *α* and stochastic perturbations.

The properties of OU are well known (e.g. Insua et al., 2012; Klebaner, 2012). The OU process is a Gaussian process with continuous paths. It has the Markov property and it is stationary, provided the initial distribution is the stationary distribution 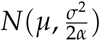. It is the *only* stochastic process which has all three properties (Gaussian, Markov, stationarity) (Breiman, 1968; Klebaner, 2012). OU is *not* a martingale. The Gaussian property of both BM and OU makes them relatively simple to work with, for example, (Butler and King, 2004; Hansen, 1997) used likelihood methods to fit models with different *μ* values on different branches of the phylogeny. Beaulieu et al. (2012) extend this idea by allowing *σ* and*α* to vary with time.

Note that the stochastic integral in (1) is with respect to “white noise”, implying that *B*(*t*) is differentiable, whereas one of the properties of BM is that it is *not* differentiable. The meaning of such integrals is therefore not straight forward. In fact, it requires a new definition for integration. The definition adopted here is that of Itô (1944, 1946). There are other approaches to stochastic integration, most notably the Stratonovich integral (e.g. Gardiner, 2009). Turelli (1977) has discussed situations in which one definition may be preferred over the other. In practice, the Ito integral is the most widely used. (Note that the ItÔ and Stratonovich stochastic differential equations for both BM and OU are identical (applying results of Gardiner, 2009, page 98).)

The main drawback of both BM and OU that I wish to highlight is the Gaussian nature of both stochastic processes. While analytically and computationally useful, this assumption limits the application of the models to trait means that are Normally-distributed across species (The distribution of traits within species is arbitrary.). Of course, one could transform the response variable so that it is then approximately Gaussian, such as using the *logit*(*x*), *probit*(*x*), or 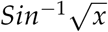 transformations for proportions, or the log(*x*) transformation for counts, and then use Gaussian process models (Ives, 2015; Warton et al., 2016). However, shoe-horning data using transformations can make interpretation of model outputs more difficult. Instead, I suggest that the direct modelling of non-Gaussian evolutionary processes provides a much more elegant view of the evolutionary process. There is a strong analogy with the development of Generalized Linear Models, which greatly extended the analysis of non-Gaussian linear models (McCullagh and Nelder, 1989). Here I outline a generalized method of constructing new stochastic process models for continuous trait evolution.

There are many more complex models that use OU as a starting point. For example, Hansen (1997) allows the evolutionary optimum to vary over time. Butler and King (2004) allow for tests of *a priori* hypotheses of different diffusion coefficients on different branches of the phylogeny. Bartoszek et al. (2012) have developed a method to analyse multivariate characters using OU. Ingram and Mahler (2013) use OU to detect evolutionary convergence in comparative data. Uyeda and Harmon (2014) use OU to analyse adaptive landscapes. Khabbazian et al. (2016) use OU to detect shifts in evolutionary optima on phylogenies. No doubt further applications will be forthcoming.

### Methods and Results

#### Diffusions as Models of Trait Evolution

Consider the stochastic differential equation (SDE):

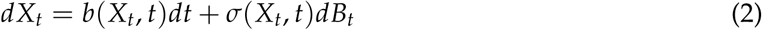

where *X_t_* is the value of our trait at time *t*. Such SDEs are termed “diffusion” equations and arise as solutions to the Fokker-Planck equation (Gardiner, 2009). The left-hand side of the equation represents a small change in trait variable *X_t_* at time *t*. The right-hand side has two terms. The first term is the deterministic part of the model. *b*(*X_t_*, *t*) is termed the *drift* function. The differential of the first term is *dt*, which denotes a differential with respect to (continuous) time. Note the difference in usage of the term compared to its use in population genetics, where drift implies a stochastic process. I will retain the traditional mathematical terminology. The second term is stochastic, as the differential is *dB*(*t*), “white noise”. *σ*(*X_t_*, *t*) is termed the *diffusion* function. In financial statistics, *σ*(*X*_*t*_, *t*) is termed the “volatility”(Mikosch, 1998). Note that both *b* and *σ* can depend on both *X_t_* and *t* in some arbitrary way. It is important that the only meaning of (2) is with respect to the Itô definition of the integral. Stochastic processes of this type are termed “Itô diffusions.”

The Ornstein-Uhlenbeck diffusion process can be defined by the following SDE:

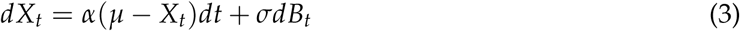

for *α* and *σ* as real, positive constants. Here, *α* represents the restraining force of stabilising selection. *μ* represents the mean trait value (at stationarity). The drift coefficient here is a linear function of *X_t_*. The form of the drift is significant, as it is this expression that controls the forcing of the trait *X_t_* back towards *μ*. OU is thus said to be “mean-reverting”: *X_t_* tends to return to *μ* over time. However, the property of mean reversion is not limited to the OU process. Eliazar and Cohen (2012) discuss the conditions where mean-reversion can occur. OU is mean-reverting largely because the Gaussian stationary distribution is unimodal and symmetric. Other processes with different diffusion functions but with the same OU drift may be mean-reverting but mode-reversion is more common, particularly in cases where the stationary distribution is asymmetric. There are cases that can be constructed where reversion is neither to the mean nor the mode. The drift function can be different to the OU drift and still be reverting to a constant, whether that corresponds to the mean or mode of the stationary distribution or another value. Hence, it is possible to reverse-engineer SDEs for any stationary distribution by judicious choice of either the drift or diffusion functions (Cai and Lin, 1996; Eliazar and Cohen, 2012).

It is also clear that (3) is time-homogeneous since neither *b* nor *σ* depend on *t*. The process is also ergodic. That is, given enought time, the time average for any particular species’ trait is equal to the average trait value across species (Lebowitz and Penrose, 1973). These properties suggest that a stationary distribution exists for this process. Figure 1 shows a sample evolution along a 5-species tree for the OU model.

**Figure. 1.**
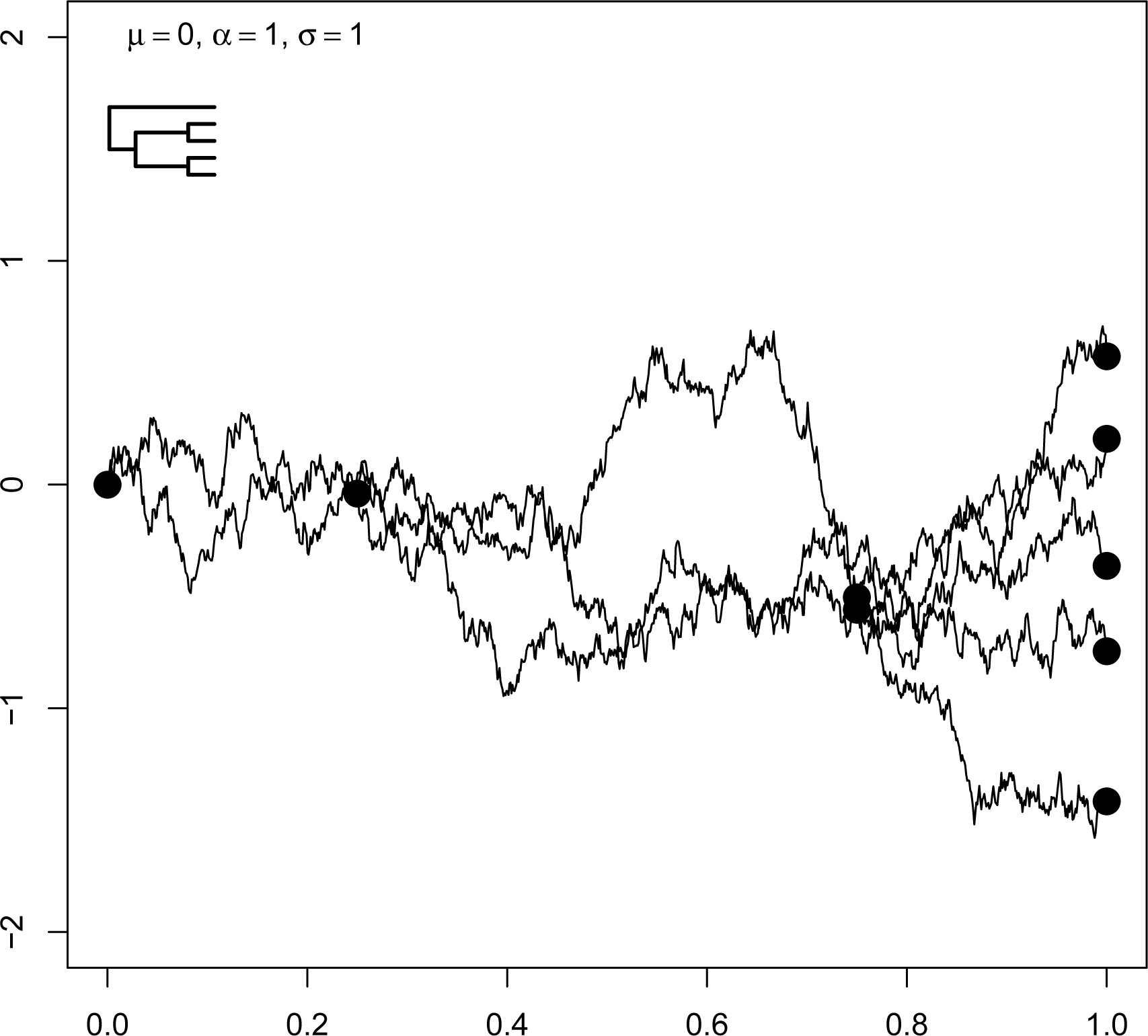
Ornstein-Uhlenbeck evolution along a 5-species tree. *μ* is the mean of the process, *α* is the strength of the restraining force, and *σ* is the diffusion coefficient. Large dots are nodes and tips.

#### New Evolutionary Models

The key to the construction of new models for evolution is the solution of the Fokker-Planck (Kolmogorov Forward) equation (Risken, 1996). In one dimension it takes the form:

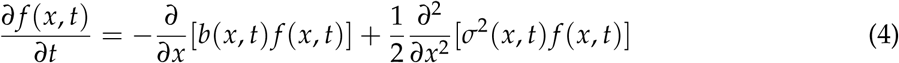

(4) governs the time evolution of the underlying probability law *f* (*x, t*). It is a partial differential equation in *x* and *t*. Note that it is *not* stochastic. If the stochastic process is time-homogeneous, that is when 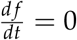, (4) can be written as:

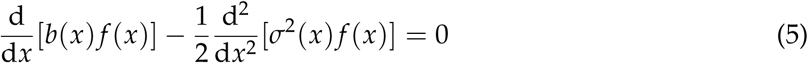

Solving for *f* (*x*) gives the following formula for the construction of the stationary distribution (Appendix 1):

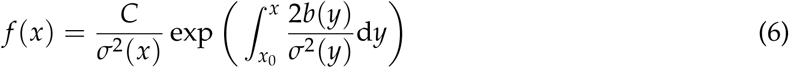

where *C* is a constant of integration found by solving *∫ f*(*x*)d*x* = 1. (6) is sometimes known as 220 Wright’s equation (Cobb, 1998; Wright, 1938).

Consider the following diffusion equations:

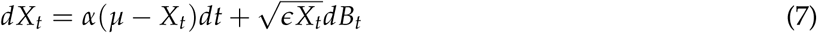

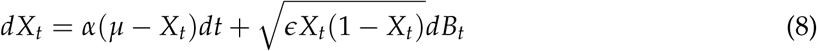

The drift terms in (7) and (8) are of the same form as in (3). Hence, these processes are both reverting, and will be driven by a central tendency towards *μ*, with a restraining force *α*. The difference between these two processes and OU is in the diffusion function. With a reverting process with OU drift, the form of the diffusion function determines the stationary distribution. The stationary distributions for each process described by (7) and (8) are derived in Appendix 2. While the notation for calculating with diffusion models is powerful and elegant, stochastic processes come alive when visualised using simulation. Example plots of paths mapped onto a phylogeny with five species are shown (Figs. 2 and 3). (7) has as its stationary distribution:

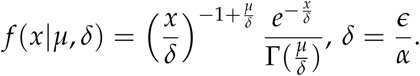

Γ is the Gamma function. That is, *f*(*x*|*y*, *μ,δ*) is a density of a Gamma distribution with mean = *μ*, mode = *μ − δ*, and variance= *δμ: x* ∼ Gamma 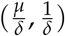. In fact, (7) is the Cox, Ingersoll and Ross (CIR) model commonly used in finance (Cox et al., 1985). See Figure 2.

**Figure. 2.**
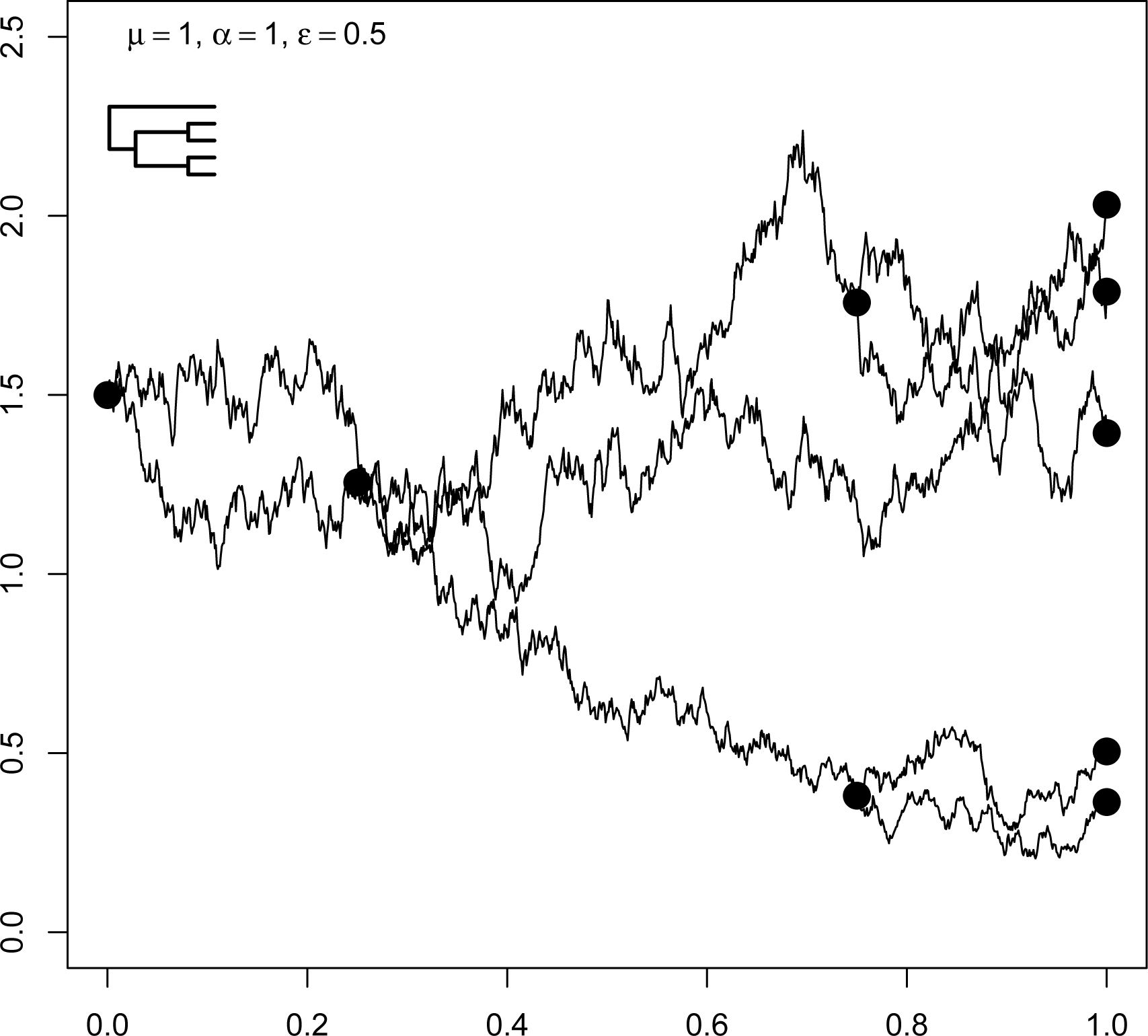
Cox-Ingersoll-Ross evolution along a 5-species tree. Large dots are nodes and tips. *μ* is the reverting level of the process, *α* is the strength of the restraining force, and *ε* is the scaling constant for the diffusion coefficient. Large dots are nodes and tips.

**Figure. 3.**
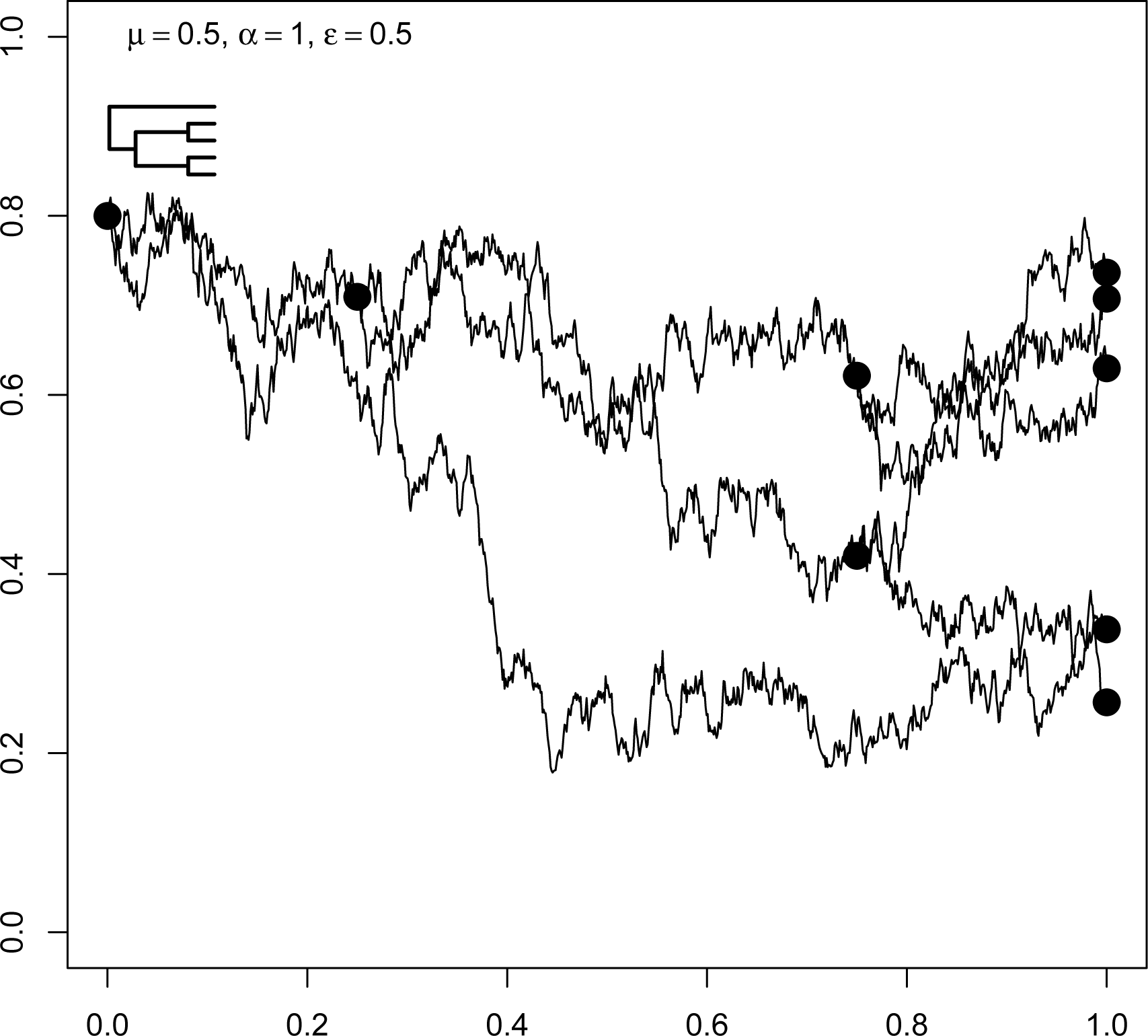
Beta evolution along a 5-species tree. *μ* is the reverting level of the process, *α* is the strength of the restraining force, and *ε* is the scaling constant for the diffusion coefficient. Large dots are nodes and tips.

The stationary distribution of the process described by (8) is:

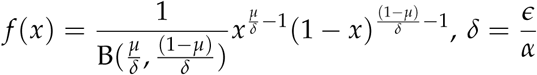

B is the Beta function. That is, *f* (*x*|*μ*, *δ*) is the density of a Beta distribution with *x* ∼ *Beta*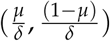

(Fig 2). The analysis of (7) and (8) and several other examples have been provided by Cobb (1998). It is interesting that in both cases, the substitution 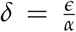 was necessary in order to correctly recognise the distributions as Gamma or Beta. This suggests that the separate estimation of *ε* and *α* is difficult if estimation is based solely on the stationary distribution. The same stationary distributions occur for arbitrary *α* and *ε*, so long as their ratio (*δ*) remains constant. A sample evolutionary path from this Beta process is presented in Figure 3.

#### Stochastic Differential Equations from Stationary Distributions

Reversing the procedure, that is deriving an SDE given a stationary distribution, is more difficult since the correspondence between SDEs and their stationary distribution (if it exists) is not unique. The problem has been addressed by Cai and Lin (1996). Extra information is needed, specifically the form of the spectral density of the process which affects the structure of the drift coefficient in the SDE. Equivalently, the autocorrelation function (ACF) of the process can be estimated and if it is absolutely integrable, the spectral density is the Fourier transform of the ACF.

Unfortunately for models of trait evolution, we rarely have detailed information on the evolutionary trajectory of a trait (ie the true historical realisation of the process) and hence we cannot analyse the spectral density of the trajectory in order to infer an appropriate drift function. If we are to proceed, we need to make extra assumptions. Fortunately, if we assume that the spectral density is of the low-pass filter type:

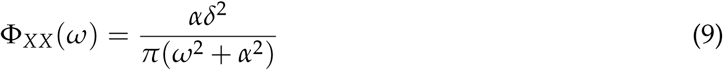

where Φ_*XX*_ is the spectral density at frequency *ω*, *δ*^2^ is the mean-square value of the process *X*(*t*), then the drift coefficient will be of the reverting OU type in (3), with *α* in (9) being identical to *α* in (3). The low-pass filter assumption implies that the drift function is determined mainly by the low frequency (long wavelength) characteristics of the evolutionary trajectory. That is, the form of the drift is mainly determined by long-lasting, slow deviations from *μ* and short-term (high-frequency) excursions are less important. To my knowledge, this assumption has never been made explicit in the literature on the application of the OU model in phylogenetic comparative methods.

Calculation of the diffusion coefficient comes directly from the application of the time-homogeneous Fokker-Planck equation (5), except instead of solving for *f* (*x*), we now solve for *σ*(*x*) (Cai and Lin, 1996). The expression for *σ*^2^(*x*) becomes:

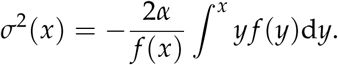

#### Transition Distributions

The stationary distribution is not the only distribution associated with a Markov diffusion process. The transition, or conditional, distribution is important for simulation and likelihood calculations (Iacus, 2008). It can be found as a solution to the Fokker-Planck equation (Klebaner, 2012) and is defined as:

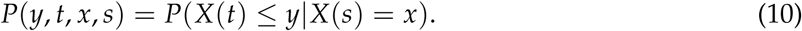

Equation (10) defines the probability distribution of *y*, the value of *X* occurring at time *t*, given that the process *X* has reached *x* at time *s*, where *s* < *t*. Unfortunately, for most processes the transition distribution is unknown or intractable. For Gaussian processes, the conditional density is usually straightforward. Brownian motion has as its transition distribution the Normal distribution with mean 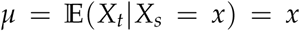, by the martingale property. The conditional variance of BM is 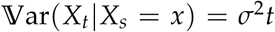. That is, the variance is independent of the trait value and depends only on *t*.

For the OU process (3) and *t* > *s* ≥ 0 the transition density is Gaussian with mean 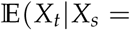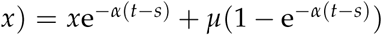 and variance 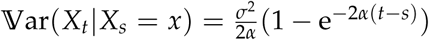. The complexity of transition distributions increases quickly with the complexity of the corresponding SDE. Equation (7), the CIR model, has the the following transition density (Cox et al., 1985):

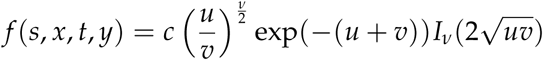

for *t* > *s* ≥ 0 where

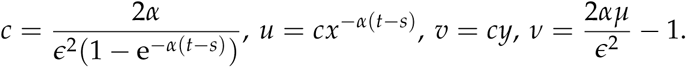

*I_v_* is the modified Bessel function of the first kind of order *v*:

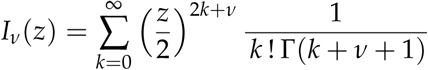

where *z* ∈ ℝ^+^ and Γ (⋅) is the Gamma function. The expectation and variance of this distribution are:

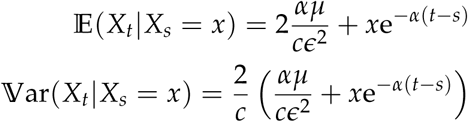

respectively. The transition density of equation (8) is even more complicated and involves infinite sums of hypergeometric functions (Abundo, 1997). Transition densities are of extreme importance for phylogenetic comparative methods, as they determine the structure of the evolutionary covariance matrix and the relationship between branch lengths and covariances (Hansen and Martins, 1996). Hence, estimation approaches such as PGLS (Blomberg et al., 2012; Grafen, 1989; Martins and Hansen, 1997) are intimately dependent on knowing transition densities. However, even quite simple models like the non-Gaussian models presented here are likely to present formidable problems with calculating evolutionary covariances, as the formulae for the transition densities for these models are extremely difficult to work with, if they are known at all.

### Discussion

#### Fitting Models to Data

To be useful, theory must be confronted with data (Hilborn and Mangel, 2013). The evolutionary models discussed here (of which BM and OU are special cases) therefore require methods to fit them to comparative data in order to estimate parameters and test hypotheses about those parameters. For BM, OU and other Gaussian processes, we can use the machinery developed for Normal distributions. In particular, there are simple relationships between branch lengths on a phylogeny and covariances for linear, Gaussian processes (Hansen and Martins, 1996). Methods have recently been proposed for the calculation of likelihoods for continuous characters on a tree, if the transition density of the evolutionary model is known (Hiscott et al., 2015). However, often the transition density is not known in closed form, or not known at all. If there was no phylogenetic dependence (that is, a star phylogeny), we could estimate model parameters, as the existence of a stationary distribution implies that at any time point independent samples will follow the stationary distribution. Unfortunately in the case of comparative data with “phylogenetic signal”, the data are not independent. We cannot make use of this result.

Simulation is currently the most popular option when the transition density of the process is intractable, although numerical solutions to SDEs are possible, particularly for large data sets (Durham and Gallant, 2002; Kloeden and Platen, 2011; Sorensen, 2004). Simulation based methods for estimating parameters for stochastic processes are widely available, largely based on theory developed for use in statistical finance (Iacus, 2008). For example, stock prices may be observed every fraction of a second, resulting in a large amount of high-frequency data with which to make inferences. Methods to deal with missing data in the high-frequency setting have been developed (e.g Roberts and Stramer, 2001). In this context the formidable problem is that data in comparative studies are only observed at the tips of the phylogeny. Rarely, internal branches may be calibrated with fossils. The entire evolutionary history of the trait for each species is thus missing and unknown. This is worse than “low-frequency” data (addressed by Fuchs, 2013): it is almost “no-frequency” data. The simulation of the entire evolutionary history, except for the tip and fossil data, is necessary. However, we may be able to combine simulated and real data using data augmentation in a Bayesian framework which might permit the approximate estimation of model parameters (Papaspiliopoulos et al., 2013; Tanner and Wong, 1987). An MCMC scheme that alternates between the update of simulated paths, and the sampling of parameters via data augmentation appears to be the most promising method (Fuchs, 2013). Such an approach would require the update of small sections (ie sub-trees) of the simulated trait history at each iteration of an MCMC procedure. Acceptance rates during MCMC are higher when only small parts of the tree are updated at a time (Elerian, 1999; Elerian et al., 2001; Kalogeropoulos, 2007; Roberts and Stramer, 2001).

The notion of using fossil phenotypes and dates to fix points in the trait-time space is attractive, but may contain grave difficulties, although recent studies have emphasised that the inclusion of fossil data can enhance our understanding of trait evolution (Slater et al., 2012). Cladistic criticisms of the use of fossils to establish ancestor-decendent relationships have never been refuted (Engelmann and Wiley, 1977; Patterson, 1981). The recent development of “tip dating” methods may avoid such criticism (O’Reilly et al., 2015; Ronquist et al., 2012). Instead we may have to be content to build quantitative trait models that incorporate ancestor-descendent relationships as ancilliary hypotheses, recognising that tests of such hypotheses may be impossible for any real dataset. However, simulation studies may be valuable in assessing the sensitivity of trait model parameter estimation to fossil placement (as an ancestor or as a sister taxon). It may be that inferring a fossil as a direct ancestor rather than as a close sister taxon will make little difference to parameter estimates for models of quantitative trait evolution. However, this has yet to be established.

The Lamperti transformation may be used to improve the simulation of trait trajectories by transforming to a unit diffusion coefficient (Burnecki et al., 1997; Fuchs, 2013; Lamperti, 1962; Moller and Madsen, 2010). Consider equation (2). The Lamperti transformation is *ϒ* = (*ϒ_t_*)_*t*≥0_ where:

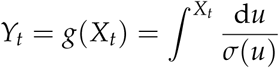

Provided the transformation g(⋅) exists and is invertible, *ϒ* fulfils the diffusion equation:

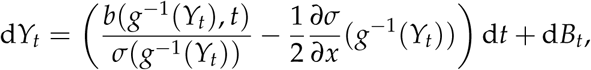

with *ϒ_t0_* = *g*(*x*_0_).

Transforming the model to remove any dependence of the diffusion coefficient on *X*(*t*) and on *t* makes the transformed process “more Gaussian” but at the cost of increasing the complexity of the drift coefficient (Iacus, 2008). However, there are grave difficulties even with fitting Gaussian models, where the transition density is known in closed form. OU has significant problems (Cooper et al., 2015), including problems with the identifiability of parameters (Ho and Ane, 2014). Many simulated likelihood methods have been proposed for fitting models where the transition density is unknown (Brandt and Santa-Clara, 2002; Cano et al., 2006; Durham and Gallant, 2002; Hurn et al., 2007; Kalogeropoulos, 2007; Sorensen, 2004), including phylogenetic comparative methods (Kutsukake and Innan, 2012). These methods often include a discretised, “locally Gaussian” approximation method such as the Euler scheme or the Milstein scheme (Elerian, 1998; Iacus, 2008). Bayesian simulation methods for parameter estimation in non-Gaussian stochastic process models of evolution is a current topic of research.

#### Stationarity

The notions of stationarity and stationary distributions have been central to this study. In the absence of an excellent fossil record of trait evolution for most traits and most taxa it seems to be a necessary, though strong assumption for evolutionary stochastic process models that are more complicated than BM. Indeed, the success of the OU process in evolutionary studies is almost as much based on its stationarity as its Gaussian properties. Several authors have constructed non-stationary evolutionary models based on OU (e.g. Bartoszek, 2012; Beaulieu et al., 2012; Jhwueng and Maroulas, 2014). Non-stationarity can arise because of time dependence in the drift coefficient, time dependence in the diffusion coefficient, or both. For mean-reverting processes, the mean of the process *μ* and/or the strength of the restraining force *α* may be time dependent (Beaulieu et al., 2012). *σ* might vary with time smoothly over the tree (Bartoszek, 2012).

Aside from the problem of overparameterisation (Bartoszek, 2012), different parameters on different clades of the tree imply at least a short period of non-stationarity as species evolve from an ancestral evolutionary regime to the new conditions. Some OU based models assume immediate stationarity after the change in evolutionary regime (e.g. Butler and King, 2004). If the old regime is almost the same as the new conditions, then stationarity in the new conditions may be achieved relatively quickly. However, if the old regime is very different from the new one, the length of the non-stationary period may be considerable and the underlying “instantaneous” stationary model will be wrong. Only fossil evidence can help in this regard because fossils can provide fixed points in the morphospace-time that can anchor the model, and provide evidence of non-stationary trait evolution or stasis. Of course, if the ancestral and derived stationary distributions are very similar so that stationarity is achieved quickly, it will be difficult to tell these two scenarios apart.

#### Model Extensions

An obvious extension of univariate stochastic processes is to re-cast them in a multivariate or multidimensional framework. There has been some research into multivariate phylogenetic comparative methods, including several software packages, largely based on BM, OU, and early-burst models (*Adams, 2014a,b,c;* Adams and Collyer, 2017; Bartoszek, 2011; Bartoszek et al., 2012; Clavel et al., 2015; Klingenberg, 2011; Klingenberg and Marugan-Lobon, 2013; Zheng et al., 2009). Certainly, multivariate diffusions are necessary to understand the correlation among characters (Bartoszek et al., 2012). However, the properties of univariate diffusion models do not always carry over to the multivariate setting. In particular, there are well-known differences between the recurrence and transience properties of Brownian motion in multiple dimensions (Morters and Peres, 2010). The analysis of the properties of multivariate diffusion models for phylogenetically-correlated data is a topic of current research (Blomberg and Rathnayake, in prep.).

A further extension of diffusion models is to the case where evolution is not strictly continuous, but is punctuated by “jumps” using Levy processes Landis et al. (2012). Levy processes are stochastic processes with independent, stationary increments. They can be thought of as consisting of three superimposed processes:

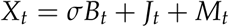

where *B_t_* is a BM (possibly with drift), *J_t_* is a compound Poisson point process, and *M_t_* is a (square-integrable) martingale with jumps. Hence, simple BM is a special case of a Levy process with no discrete jumps. Note that OU is not a Levy process. Landis et al. (2012) estimate parameters for a Levy process fitted to data for body mass and brain volume in primates, and found evidence for some jumps in each trait, rejecting a simple BM model. Duchen et al. (2017) used Levy process models to look for jumps in *Anolis* lizard body size and Australasian lories (Aves: Loriini) gut morphology. Both taxa showed evidence for evolutionary jumps. The application of Levy processes to phylogenetic comparative data is promising, but given the difficulties and complexities of fitting Itô diffusions, it may pay to be wary of hidden pitfalls. Certainly the *post hoc* identification of jumps may not be o9f much use without a working hypothesis for *why* we may expect jumps at certain nodes or on certain branches on the tree, and models already exist for postulating *a priori* different rates of evolution in different parts of the tree (e.g. Butler and King, 2004; O’Meara et al., 2006). It may be difficult to choose between “jump” models and models that estimate rapid changes of evolutionary rate (large differences in *σ*) for particular clades (e.g. Alfaro et al., 2009; Rabosky et al., 2014, 2013; Shi and Rabosky, 2015), although jump models tend to have fewer parameters, which may make them more parsimonious.

Models with jumps may be a good approximation in situations where evolution, though continuous, has been so rapid that on the macroevolutionary timescale change appears to be instantaneous (Duchen et al., 2017). Landis and Schraiber (2017) find evidence for “jumps” in the evolution of vertebrate body size and interpret this as rapid evolution, followed by stasis.

#### Evolutionary models for phylogenetic comparative analyses

Scientific models may be developed with several different motivations (Gavrilets, 1999). The scientist may build models to make a decision (e.g. to reject a null hypothesis), summarise evidence (e.g. calculate the likelihood of observing the data, given a model) or quantify their beliefs (using Bayes Theorem). Another important property of a model is its predictive ability, and predictive models have long been the favourite approach in the physical sciences: models predict future observations which then test the validity of the model. Biologists, and especially evolutionary biologists, have never put much faith in predictive models (Gavrilets, 1999; Hillis, 1993). So many factors affect the evolution of organisms, and over such a long timespan, that one is tempted to give up hope of developing mathematical models that have any predictive value in the real world. And it is true that it would be foolish to make predictions of where in the phylomorphospace (*sensu* Sidlauskas, 2008) species will evolve to in some future deep time. We have no hope of making the necessary observations.

Although we may not be able to predict the precise evolutionary trajectory of any particular species, we can perhaps predict (or postdict) the probability distribution of traits across species. We may use crossvalidation (Efron and Gong, 1983) to assess the predicitve ability of our models, or in a Bayesian context posterior predictive simulation (Gelman et al., 2013). Given the traits from a newly discovered species (fossil or extant), we can predict that the new trait values fit well within the distribution of the known species’ trait values, where the parameters of the distribution are estimated from extant species using a particular model of evolution. If the values for the new species’ traits are more extreme so that they fall into the tails of the stationary distribution, we may reject our model of evolution for that set of species and traits. Predictions of the models may not be precise, but they are predictions nonetheless (see Pigliucci, 2007). This “grey box” approach to model identification (Kristensen et al., 2004) gives up the possibility of knowledge of the microevolutionary processes leading to species diversification and trait evolution. This is replaced with a tractable stochastic process that summarises the evolution of the statistical distribution of trait values over deep time. Given the poor quality of many comparative data sets(little or no fossil data, incomplete sampling of extant taxa, low intraspecific replication), this may be the best that can be achieved.

The modelling approach and the new models described here involve a considerable amount of mathematical sophistication in their derivation and in the analysis of their properties. Computational skill is necessary in developing algorithms to fit the models to data. Critics may object that the approach outlined here is too complex or unnecessary. However, diffusions are already the most popular model for phylogenetic comparative studies, in the form of BM and OU. The present author hopes simply to widen horizons and provide a unifying framework. It is true that all models are wrong but some are useful (Box, 1976). Nevertheless, mathematics (and its sister taxon, computation) are the best tools we have in order to precisely describe both the nature of macroevolutionary phenomena and our assumptions about them. A small amount of precise mathematics can sometimes cut through imprecise verbal arguments. For example, the microevolutionary genetic theory developed by Fisher, Haldane and Wright effectively silenced the arguments between naturalists and Mendelians on the importance of natural selection and the nature of genetic variation, leading to the Evolutionary Synthesis (Mayr and Provine, 1998). A mathematical theory of macroevolution which unites stochastic models of trait evolution with models of phylogenesis, speciation and extinction may allow us to better statistically model the course of phenotypic evolution (e.g. FitzJohn, 2010; Goldberg et al., 2011; Maddison et al., 2007), although estimating parameters for these models may be difficult without fossil trait data. Recent applications of trait-mediated diversification models based only on extant trait data may be misleading (Rabosky and Goldberg, 2015). A more sophisticated understanding of the mathematics of diffusions and other stochastic processes may allow the critical appraisal of macroevolutionary models for biological phenomena in deep time.

## Conclusion

Currently, popular models of trait evolution rely heavily on Gaussian processes and their useful mathematical properties. However, non-Gaussian models are possible and may have some advantages over Gaussian models in certain situations where the data are likely to be non-Normal. The present study describes new, non-Gaussian models of trait evolution, together with methods for building new models, and a discussion of the mathematical and computational difficulties in working with diffusion models in a more generalised setting. Several new avenues for investigation are suggested. In particular, the role of fossils in improving the identifiability of models and the extension of models to multivariate trait space seem especially timely. These areas are not without challenges. Including fossils as ancestors, rather than as sister taxa has been a difficult problem for many years, as the early cladists were well aware. The extension of univariate models to multivariate trait space is likely to be more difficult than expected, as even the simplest evolutionary model, BM, has different properties in multiple dimensions. Another important research direction is to establish the expected covariances for traits in terms of the transition distributions for non-Gaussian models. This is likely to be difficult but would pay off immensely, allowing estimation of regression models by Phylogenetic Generalised Least Squares (PGLS), allowing the construction of a new Generalized Phylogenetic Model, by analogy with Generalized Linear Models. Nevertheless, research into the application of stochastic process (diffusion) models to the evolution of quantitative traits appears to hold great promise.

## Appendix 1: Derivation of Wright’s Equation

Consider the Fokker-Planck equation for an Itô diffusion *X*_*t*_ (4). Alternatively, (4) can be rewritten as (Risken, 1996):

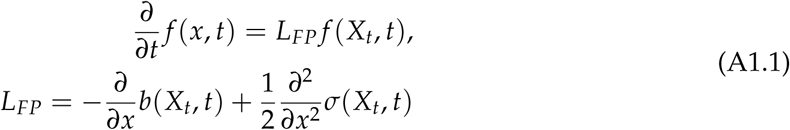

Further, equations (A1.1) can be written as:

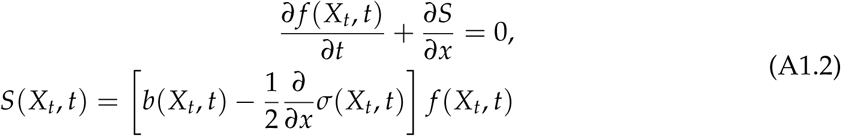

*S*(*X*_*t*_, *t*) can be interpreted as a probability flow. For natural boundary conditions min *x* = − ∞ and max *x* = ∞, and assuming time-homogeneity, *S*(*X_t_*, *t*) = *S*(*X*_*t*_) = 0. Letting *x* = *X*_*t*_ we have the following first-order linear differential equation:

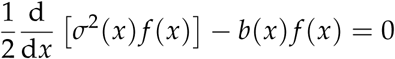

Let *m*(*x*) = *σ*^2^(*x*)*f* (*x*), implying 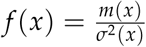 then

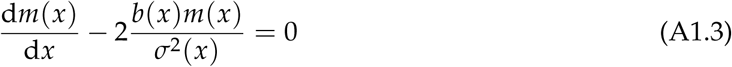

Equation (A1.3) can be solved using the method of integrating factors. Let 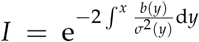.

Multiplying both sides of equation (A1.3) by *I*:

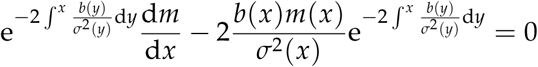

Integrating both sides and using the product rule on the LHS,

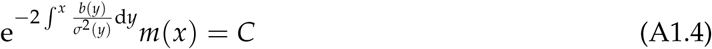

where *C* is a constant of integration. Substituting *m*(*x*) = *σ*^2^(*x*)*f*(*x*) and rearranging, we have:

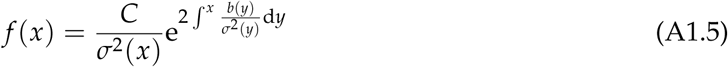

which is Wright’s formula.

## Appendix 2: Derivation of Stationary Distributions

**CIR model**

Let 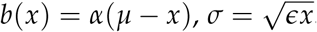. Substituting into Wright’s formula (6):

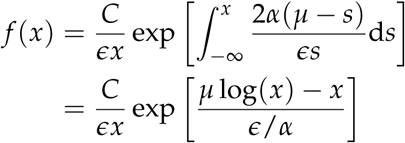

Let 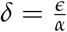. Then:

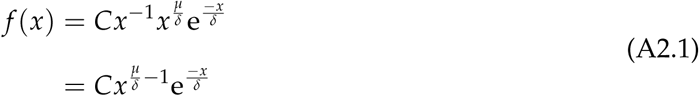

Equation (A2.1) can be recognised as the kernel of a Gamma density, with shape 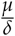 and scale *δ*, and with normalising constant *C*. Therefore,

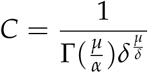

where Γ(⋅) is the Gamma function. i.e.

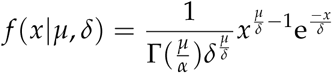

or *x*|*μ*, *δ* ∼ Gamma 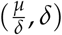.

**Beta model**

Let 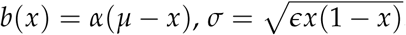. Substituting into Wright’s formula (6):

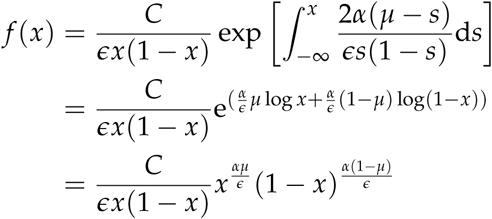

Setting 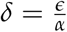 and simplifying further, we have:

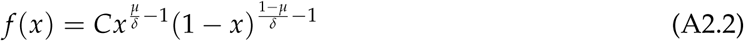

Equation (A2.2) is the kernel of a Beta distribution with shape parameters 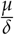 and 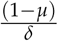 and normalising constant *C*. Hence, the density can be written as:

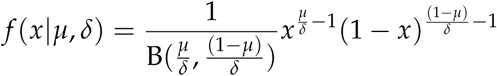

where B(⋅,⋅) is the Beta function. More succinctly, *x*|*μ*, *δ* ∼ Beta 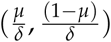

